# Oxidation-induced structural changes in actin and myosin evaluated by computational simulation, machine learning modeling and high-speed AFM

**DOI:** 10.1101/2024.07.04.600628

**Authors:** Oleg S. Matusovsky, Daren Elkrief, Yu-Shu Cheng, Dilson E. Rassier

## Abstract

High levels of reactive oxygen species produced during muscle oxidative stress are implicated in the development of several muscle diseases. To better understand the mechanism behind a reduced myosin force generation under oxidizing conditions, we analyzed the structural and functional changes in the actin and actin-myosin complex using high-speed atomic force microscopy (HS-AFM), simulated HS-AFM, and molecular dynamics (MD) simulation. Computational oxidative nitration of tyrosine residues demonstrated instability in the molecular structure of the F-actin subunit. Cross-section analysis of the simulated HS-AFM images revealed a shift in the height values (∼0.2-1.5 nm in magnitude) between the non-oxidized and oxidized actin, which correspond to the height differences observed in HS-AFM experiments with in vitro oxidized F-actin. The oxidation-induced structural alterations in actin impact myosin molecule displacement on the single-molecule level. The displacements of myosin heads along the F-actin filaments in the presence of ATP involve the binding of the myosin molecule to a specific site on the F-actin filament, followed by the rotation of the myosin lever arm, which triggers the release of inorganic phosphate (Pi). Subsequently, the myosin head detaches from the F-actin and re-binds to a new site on the filament. The formation of the SIN-1-treated F-actin-myosin complex in the presence of ATP resulted in a change in myosin head displacement size, with a significant decrease in the frequency of long displacements (≥ 4 nm). These results suggest that oxidation decreases the pool of the weak-bound myosin molecules and shortens the long displacements related to the Pi release step, reducing the force generation by myosin motors.

## Introduction

Oxidation is the result of an imbalance between production of reactive oxygen / nitrogen species (RONS) and an elaborate antioxidant defense system. The process causes an “oxidative damage” to different cell systems, including muscle, leading to various pathological conditions: muscular dystrophy**^1^**, muscle weakness in rheumatoid arthritis**^2^**, sarcopenia or age-related muscle loss**^3–5^**, skeletal, cardiac myopathies and heart failure**^6–8^**. These manifestations can be attributed to the significant impact of oxidation on protein structure. Proteins can undergo oxidative modifications that alter their physical properties, including changes in conformation, structure, solubility, susceptibility to proteolysis, enzyme activities and interactions with other molecules.

In skeletal muscle, RONS such as hydrogen peroxide (H_2_O_2_), nitrogen oxide (NO^●^), peroxynitrite (ONOO^●-^) or hydroxyl radical (^●^OH) have been shown to directly alter contractile function**^9^**. The exact mechanism of the impaired contractility upon oxidation is still not clear, since the susceptibility of proteins to oxidation varies depending on the specific amino acids involved in the process**^10^**. Thus, a broad range of manifestations due to protein oxidation has been observed *in vitro*: reducing motility of actin filaments in the *in-vitro* motility assays**^11,12^**, inhibition of the rate of polymerization of actin**^2^** and the myosin ATPase activity**^13^**, reducing force generation in the muscle fibers, myofibrils and myosin filaments**^12,14,15^**. It was suggested that oxidation may lead to a reduction in the number of myosin cross-bridges and thus a decrease in force generation**^15^** or increase in the rate of myosin head detachment from actin, further reducing muscle force. Exposure of actin and myosin to oxidative agents affects actin-myosin interactions, leading to a decreased myosin-propelled actin velocity and a reduction in force generated by myosin and actin filaments**^12^**. Moreover, oxidative modifications on actin alone were sufficient to affect myosin force and velocity on actin filaments**^2,12,13^**. Nevertheless, the underlying mechanism of the impaired ability of myosin cross-bridges to generate force at the molecular level is still poorly understood with multiple factors yet to be identified.

The purpose of this study is to identify the myosin force generation steps under oxidation conditions at the single-molecule level and high spatiotemporal resolution. We tested the idea that the myosin functionality is disrupted due to structural changes caused by the accumulation of oxidative post-translational modifications in actin filaments and/or in myosin heads. Using high-speed atomic force microscopy (HS-AFM) we showed that oxidative modifications lead to structural changes in the skeletal muscle myosin N-terminal domain and actin filaments. The oxidized myosin molecule displayed a notable change in the spatial orientation of the myosin heads compared to the non-oxidized molecules. This altered head orientation impacted the binding capacity of the oxidized myosin to non-oxidized actin filaments in our HS-AFM experiments. Conversely, the oxidized actin filaments, when combined with non-oxidized myosin in the presence of ATP, led to a reduction in the size of myosin head displacement and force generation. Furthermore, the structural impact of oxidative nitration on actin subunits was investigated using molecular dynamics (MD) simulation and simulated HS-AFM images aimed to uncover the functional consequences of this modification on actin’s structure.

## Results

### Dynamics of actin-myosin interaction under oxidation conditions

To study the functional activity of skeletal myosin in the actin-myosin complex under oxidation conditions, we first tested the situation when the heavy meromyosin (HMM), a double-headed proteolytic fragment of myosin, is oxidized by 5-amino-3-(4-morpholinyl)-1,2,3-oxadiazolium chloride (SIN-1). SIN-1 can generate both nitric oxide (NO) and superoxide (O2-) in aqueous solutions. These two species then react to form peroxynitrite (ONOO-), a potent oxidant and nitrating agent. Therefore, SIN-1 is often used to study the effects of oxidative stress on biological systems. However, in our HS-AFM experiments, the frequency of successful interaction between the SIN-1-treated HMM and F-actin attached to the mica-supported lipid bilayer (mica-SBL) surface was notably low. One of the possible reasons can be attributed to the structural characteristics of the SIN-1 oxidized HMM investigated in dynamics. As shown in Figure 1a, the oxidized SIN-1 HMM exhibited a significant difference in the spatial orientation of the myosin heads from the non-oxidized HMM molecules. The oxidized heads appeared to be interlinked with the progression of the linkage over time (Movie S1). Indeed, the distance between two myosin heads within one molecule of the SIN-1-treated HMM was significantly reduced as observed from the Gaussian distributions (Fig. 1b).

**Fig. 1.**
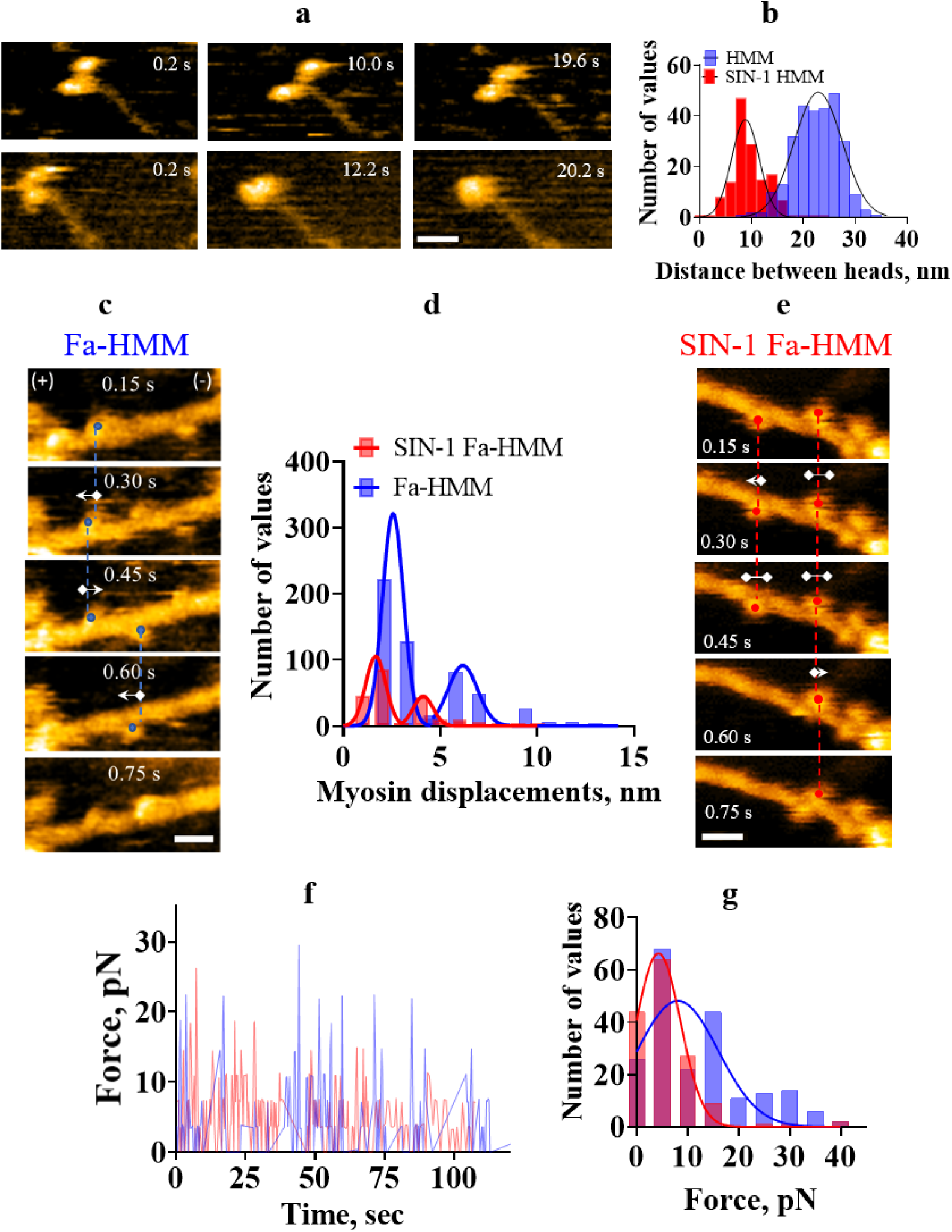
Oxidation-induced structural changes in HMM molecules and the HMM displacement. **(a)** The non-oxidized and SIN-1-treated skeletal HMM, a double-headed fragment of myosin, on the mica-APTES surface. Scan area: 150 × 75 nm^2^, 80 × 40 pixels^2^, temporal resolution: 5 frame per second, Scale bar: 30 nm. **(b)** The distance between two HMM heads measured over time: 8.9 nm ± 2.7 nm (SD) for SIN-1-treated HMM and 22.9 ± 4.6 nm (SD) for non-oxidized HMM. **(c, e)** HS-AFM images of skeletal HMM bound to the non-oxidized F-actin **(c)** and SIN-1 treated F-actin (**e**) in the presence of ATP. Scan area: 150 × 75 nm^2^, 80 × 40 pixels^2^, temporal resolution: 6.7 frame per second. Scale bar: 30 nm. Arrows with symbols, 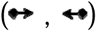: forward or backward head displacement between reference frame and subsequent frame; 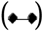: no head displacements between reference frame and subsequent frame. **(d)** Histogram of the HMM displacements along the non-oxidized F-actin (blue) and SIN-1 treated F-actin (red). Data were fitted by two Gaussians: r^2^=0.99, r2=0.90 and r^2^=0.99, 0.91 for Fa-HMM and SIN-1-Fa-HMM, respectively; n=758 events (Fa-HMM), n=303 events (SIN-1-Fa-HMM). The average displacements for Fa-HMM: 2.6 nm ± 0.5 nm (1^st^ peak) and 6.1 nm ± 0.8 nm (2^nd^ peak); for SIN-1-Fa-HMM: 1.7 nm ± 0.5 nm and 4.0 ± 0.5 nm (2^nd^ peak) **(f, g)** The force produced by myosin molecule in the Fa-HMM (blue) and in the SIN-1-Fa-HMM complex (red) calculated from myosin displacements over time (**f**) and plotted as a histogram (**g**). The force values were calculated from the equation *F = k × d*, where *k* is a stiffness of HMM equal to 2 pN/nm per head**^18^** and *d* is a displacement of myosin molecule along the actin filament observed in the present study.

Due to the impact of SIN-1 oxidation on the myosin heads orientation, resulting in the weak binding of SIN-1 oxidized HMM to F-actin in our HS-AFM experiments, we chose to utilize SIN-1 treated F-actin complexed with non-oxidized HMM. This approach allowed us to explore the effects of SIN-1 oxidation on the actin-myosin complex in a more controlled manner, providing insights into the structural and functional alterations of the myosin power stroke at the single-molecule level in dynamics.

After attachment of oxidized or non-oxidized actin filaments, myosin HMM molecules were added to the HS-AFM camber to form the actin-myosin complex in the presence of ATP. During the power strokes, the HMM molecules interact with either non-oxidized or SIN-1 oxidized F-actin and move myosin heads forward or backward along the actin filament (Movie S2). The analysis of the myosin displacements is detailed in our previously published studies**^16,17^** and briefly outlined in the Material and Methods.

### The analysis of myosin displacement and force generation in HMM-F-actin complex

The HMM heads displaced on the actin filaments as a consequence of the force-generation steps and cycling interaction with actin filaments in the presence of ATP. In terms of myosin cycling and the power stroke we defined the changing of the HMM position through a detachment and reattachment to the new binding site along F-actin as a transition from the weak to the strong-binding states that occurred within one or a few frames. In the presence of ATP, myosin heads detach from the actin filament and re-attach to the same or a new binding site, allowing us to determine the displacement by the evaluating the change in the averaged center of mass of the myosin heads between the reference frame and the next frame in HS-AFM successive images. For instance, if a myosin head binding to F-actin in one frame was observed in a different position in the next frame, it was considered as being detached and reattached to a new binding site. On the other hand, if the myosin head remained in the same position on F-actin for several frames without any change in its center of mass, it was assumed there were no displacement.

Typically, under our experimental conditions (low ionic strength with 25-100 mM KCl, 2-4 mM MgCL_2_, 1 mM EGTA and neutral pH 7.0), the displacement distribution of the myosin heads bound to the non-oxidized F-actin showed three common sizes: short displacements (under 1 nm), moderate displacements (∼1-3 nm) and long displacements (over 3-4 nm). According to various experimental data, the long displacements ranged from 3-4 nm and over with averaged values ∼4.0-6.1 nm, as in our study, most likely represent the Pi release step**^16,19,23^** and lever arm movements that are responsible for force-generating step. The short displacements ranged from 0.2-2.9 nm with averaged values ∼1.7-2.6 nm most likely represent the ADP release step**^19–22^**. This pattern of displacement distribution was also observed in our previous studies, where non-oxidized actin-myosin complex was used**^16,17^** and showed similar displacement size with experimental data for cardiac and skeletal myosins**^20,23^**.

Comparably to the previous published data, the displacement distribution of HMM heads along the F-actin was best fitted by sum of two Gaussians function and showed two main peaks. The average displacements for non-oxidized Fa-HMM complex: 2.6 nm ± 0.5 nm (1st peak) and 6.1 nm ± 0.8 nm (2nd peak); for SIN-1-Fa-HMM complex: 1.7 nm ± 0.5 nm and 4.0 ± 0.5 nm (2nd peak) (Fig. 1d). The displacement distribution of the HMM molecules complexed with SIN-1-treated F-actin revealed the shift to shorter displacements, suggesting that the frequency of long displacement events was significantly decreased by the oxidation (Fig. 1d). The force generated by the HMM molecules in the non oxidized Fa-HMM and the SIN-1-Fa-HMM complexes was determined from the myosin displacements over time (Fig. 1f) and presented as a histogram (Fig. 1g). The force values were calculated using the equation F = k × d, where k represents the stiffness of HMM and d is the displacement of the myosin molecule along the actin filament, as observed in the present study. Previous studies utilizing different models and experimental approaches, such as single molecule and muscle fibers measurements, have determined the stiffness of a myosin molecule or HMM to be approximately 2 pN/nm per head **^18,19,24–26^**.

### HS-AFM visualization of the actin filaments treated by SIN-1

The obtained data suggests that the impact of oxidation on myosin’s functional activity can be either direct or indirect. The direct effect refers to structural changes in the SIN-1-treated HMM heads, which appeared to be interconnected under oxidation conditions (Fig. 1, Movie S1). These structural alterations make it impossible to visualize SIN-1-oxidized HMM binding to F-actin in our HS-AFM experiments. On the other hand, an indirect functional outcome was observed when the actin-myosin complex was formed with SIN-1-treated actin filaments and non-oxidized HMM. This finding is particularly intriguing because it allowed the observation of the functional outcome of the myosin molecule based on potential structural changes in the actin filaments, prompting a more detailed investigation of this effect.

We conducted HS-AFM experiments on both untreated and SIN-1-treated skeletal F-actin. The SIN-1 treatment was performed according to our previous studies**^12^** and described in the Methods. The imaging of the non-oxidized F-actin and the oxidized by SIN-1 F-actin attached to the mica-supported lipid bilayer surface (mica-SLB) was performed by the same cantilever to avoid any difference caused by tip cantilever collision during HS-AFM scanning (Fig. 2b). There were no differences observed for the half helical pitch distances between the non-treated and SIN-1-treated F-actin (Fig. 2c). Conversely, the height distributions measurements of the highest points of the peak at the cross-over of the double F-actin helix, revealed significant difference between the non-treated and SIN-1-treated F-actin: 8.5 nm ± 2.9 nm (SD) and 7.0 nm ± 1.2 nm (SD) (P=0.006), respectively (Fig. 2d).

**Fig. 2.**
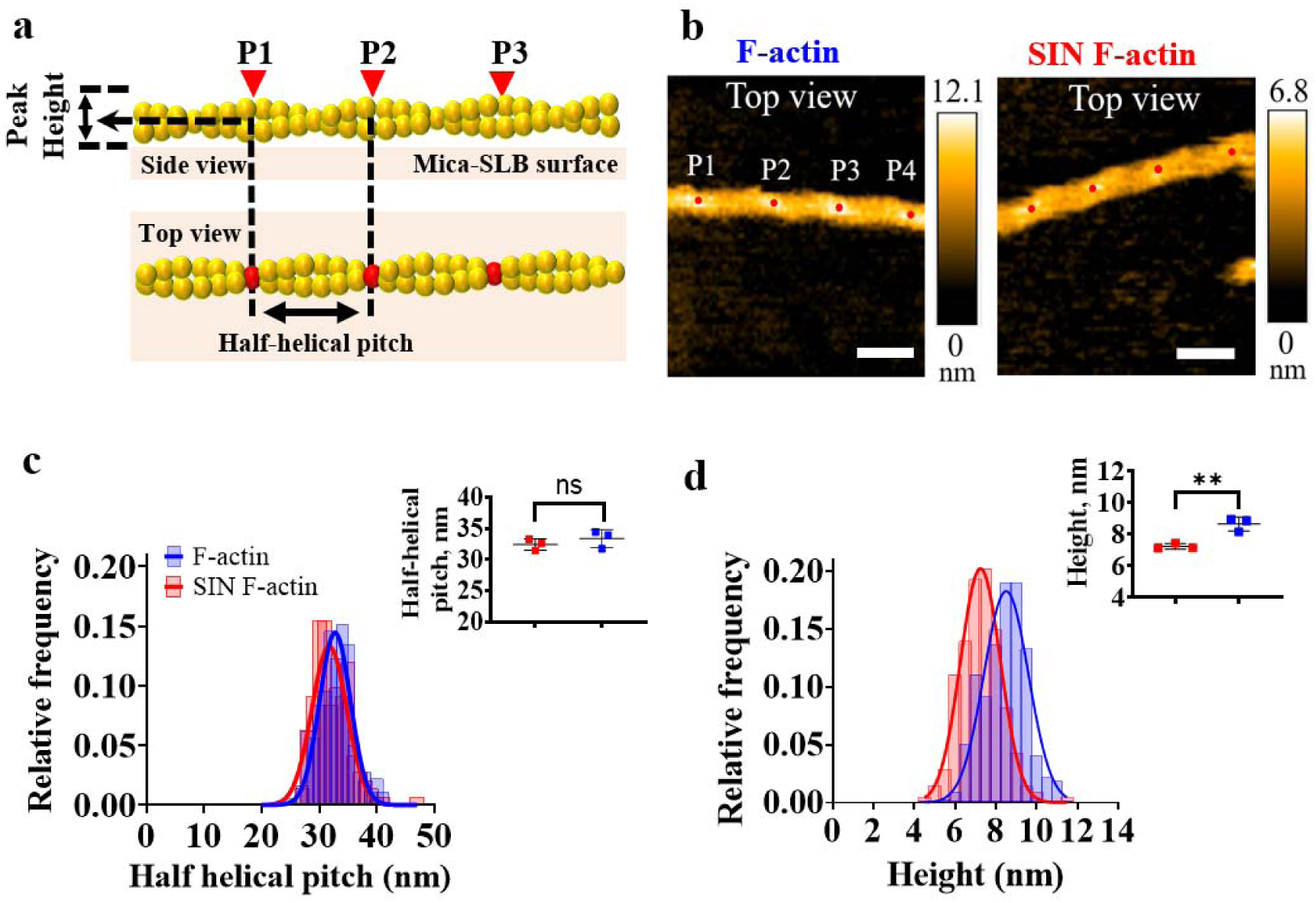
Comparison of the height profiles of the actin filaments treated by SIN-1 in the experimental and simulated HS-AFM images. (**a**) Diagram representing the peak height measurements along actin filament. The red points show the peaks (P) in the actin structure with the highest values as a cross-over of the double F-actin helix. **(b)** Representative HS-AFM images of control and SIN-1-treated actin filaments located on mica-supported lipid bilayer with corresponding Z-scales in nm. Scan area: 150 x 150 nm^2^ (80 x 80 pixels^2^), scale bar: 35 nm. **(c, d)** Peak height and half-helical pitch measurements for the non-treated and SIN-1-treated F-actin. No differences were noted in half-helical pitch between control (n = 178) and SIN-1 filaments (n = 142, P = 0.3994), whereas peak height (d) was significantly lower in SIN-1 filaments (n = 207) compared to controls (n = 315, P = 0.006). All values shown as mean ± SD.

### Computational analysis of the non-treated and 3-NT oxidized G-actin structure

The observed differences in height values between oxidized and non-oxidized F-actin indicate that SIN-1 treatment induces structural changes in actin filaments. As mentioned, SIN-1 can produce both NO and O2-in aqueous solutions, which form peroxynitrite (ONOO-), a potent oxidant and nitrating agent. Indeed, the consistent occurrence of 3-nitrotyrosine (3-NT, NO2; +46 Da) and malonaldehyde (MDA, C3H3O; +54 Da) modifications in myofibril actin filaments and purified skeletal muscle F-actin were identified by mass-spectrometry**^2,27–28^**. Furthermore, it was shown that the nitration of tyrosine (Tyr) residues, such as Tyr53 (near D-loop), Tyr91, Tyr198, and Tyr240 in F-actin impair enzymatic function, leading to alterations in actin polymerization, stability of actin filaments, actin-myosin interaction and myosin ATPase activity**^2,13, 29^**. After considering these results and our observations, we decided to study the impact of tyrosine nitration (3-NT) on the structures of G-actin and F-actin in more detail.

To initiate computational assessment and to make a connection with experimental data, certain tyrosine residues in the F-actin model subunit (PDB 2ZWH, 3.3 Å) or F-actin (PDB 6BNO, 5.5 Å) structures were subjected to the oxidative nitration (3-NT) in the PyMOL environment (v.2.3.3, Schrodinger LLC) by adding the NO_2_ group to the phenyl ring of the tyrosine. The tyrosine residues that we modified were identified by mass-spectrometry analysis in actin filaments of patients with rheumatoid arthritis**^2^**. The *in-vitro* treatment of purified skeletal G-actin and myofibril F-actin with SIN-1 also caused an oxidation of similar tyrosine residues**^2^**. The following residues in the G-actin subdomains (SD) were modified: Tyr 91, Tyr 362 (SD1); Tyr 53 (SD2); Tyr 166, Tyr 169, Tyr 294, Tyr 296, Tyr 297, Tyr 306 (SD3); Tyr 188, Tyr 198, Tyr 218, Tyr 240 (SD4) (Fig.3, Movie S3, Movie S4).

**Fig. 3.**
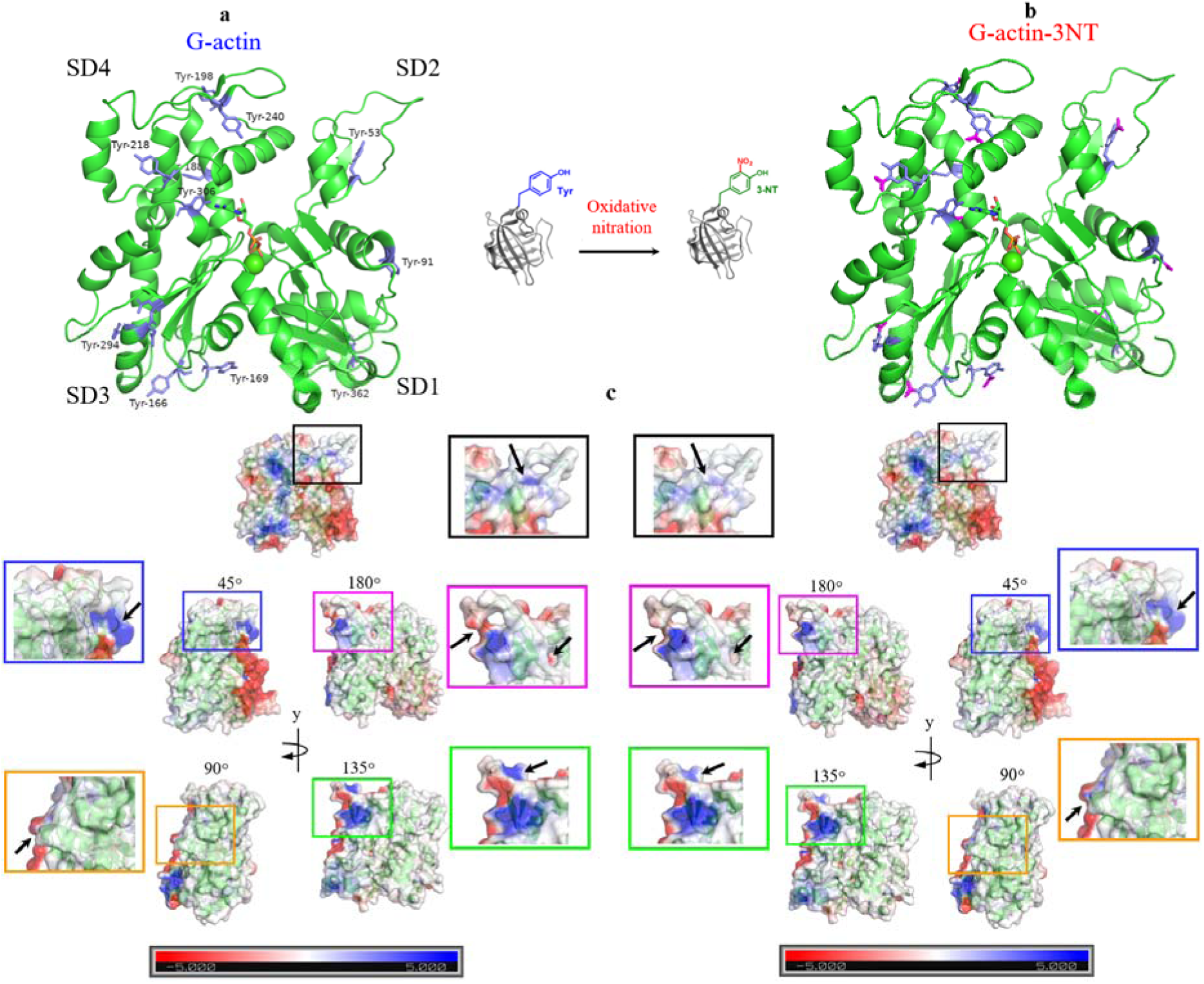
Computational oxidative nitration of the G-actin. **(a)** The non-oxidized and **(b)** computationally oxidized G-actin (PDB 2ZWH). The modified residues shown as blue sticks with the phenyl ring in each subdomain (SD1-SD4). Oxidative modifications were performed in the pyMOL using pyTM plugin (v.1.2)**^30^**. The oxidized atoms in the residues highlighted by pink color. **(c)** The successive y-axis rotation of the non-oxidized and oxidized G-actin structures with 45° step from in tial position showed the local changes in the electrostatic potential energy around oxidized residues. The matched colored boxes for each orientation were magnified to show the charge differences more precisely. The map scale represents the electrostatic potential energy from −5.0 k_B_T (red) to 5.0 k_B_T (blue).

The comparison of the G-actin electrostatic maps revealed that the total charge of the G-actin monomer was not affected by the presence of the residues modified by the oxidative nitration: −12.99 for both G-actin and G-actin-3NT. Similarly, the electrostatic potential (U) between two arbitrary charges q1, q2 separated by distance r was calculated as the sum of the potential energy of all pairs of atoms in the structure divided by the distance between the charged atoms, according to Coulomb’s law: U = q1* q2 / r2 was not different in both structures. However, the successive rotation of the non-oxidized and the oxidized structures around the y-axis (Fig. 3) or x-axis (Fig. S1, Movie S5) revealed several spots affected by oxidative nitration. According to the electrostatic map, the spots were recognized as a change of the positive / negative charges (blue and red colors, respectively) in the non-oxidized structure to the neutral charge in the oxidized structure.

### Molecular dynamics simulation

To test whether the observed local changes in the electrostatic charges around Tyr-3NT residues affect the protein conformational dynamics in silico, we performed the molecular dynamics (MD) simula ion of G-actin monomer with non-oxidized and Tyr-3NT oxidized residues. The details for MD simulation described in the Methods. The custom Python code, based on the OpenMM package**^31^**, was utilized for MD simulation can be accessed on GitHub: https://github.com/matusoff/Molecular-dynamics-with-BioPython and in the Supplementary Data. The OpenMM package provides a versatile set of libraries for molecular simulation, enabling the integration of various force fields and the execution of MD simulations. Additionally, the MDAnalysis library offers robust tools for the analysis of MD trajectories, further enhancing the capabilities for in-depth trajectory analysis.

Running the MD simulation for 100 ns at the temperature of 300 K and subsequent analysis of the resulting trajectory files revealed that the equilibration of G-actin structure, as judged by a root-mean-square deviation (RMSD), was happened after ∼20 ns of the simulation. Therefore, this conformation of the G-actin monomer can be used as the favoured conformation at the pH 7.0, 300 K and ionic strength 0.15 M (Movie S6, Movie S7). In the text below, this conformation will be titled as the in tial conformation. In contrast, G-actin structure with Tyr-3NT residues showed a less stable behaviour with the areas of increasing the RMSD approximately every 10 ns (Fig. 4c). Thus, MD simulation demonstrated that 3-NT oxidative nitration of the tyrosine residues in G-actin contributes to the instability of the system.

**Fig. 4.**
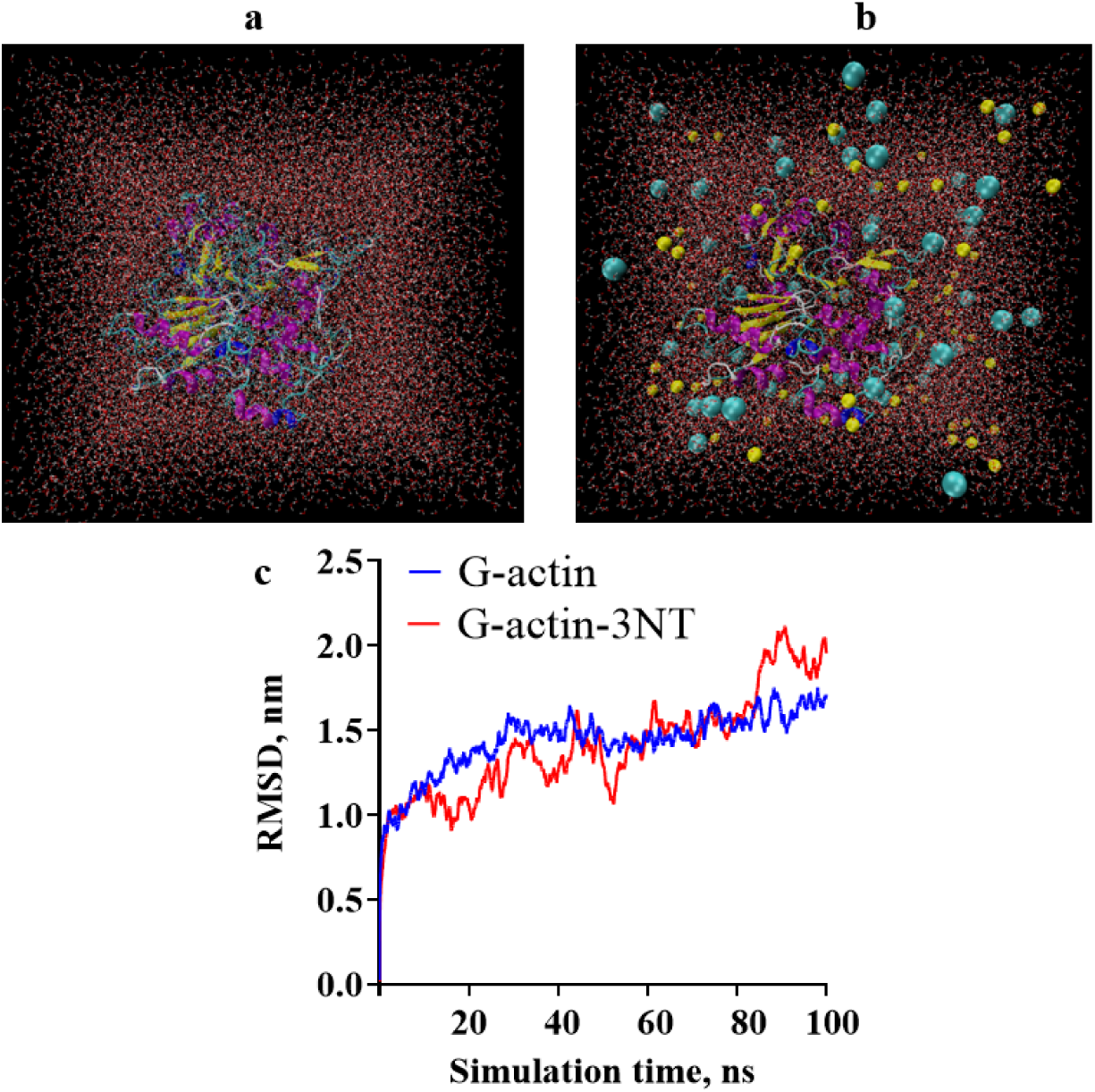
Molecular dynamics (MD) simulation of the G-actin structure with non-oxidized and Tyr-3NT oxidized residues. An example of solvation **(a)** and ionization **(b)** of the G-actin structure. The structure was placed in a rectangular box filled with water molecules. The cyan and yellow spheres represent Na^2+^ and Cl^-^ ions, respectively. **(c)** The root-mean-square deviation (RMSD) calculated for G-actin structures with non-oxidized and Tyr-3NT oxidized residues.

### HS-AFM simulation of the G-actin / G-actin-3NT structures

To further study the conformational changes in the area around oxidized residues we simulated HS-AFM image with a BioAFM viewer (v.2.5, Kanazawa University), a recently developed open-source software**^32,33^** that simulates HS-AFM images from the PDB files of the atomic structure with high correlation (Fig. S3). The G-actin structures with the non-oxidized Tyr and the oxidized Tyr-3NT residues were used to simulate the HS-AFM images of the non-oxidized and the oxidized G-actin structures in the initial and rotated conformations (Fig. 3, Fig. S2, Movie S5). The BioAFM viewer allowed to change the apex angle and tip radius to obtain the HS-AFM simulated images close to experimental conditions (Fig. S4).

The simulated HS-AFM images were used to gain the height profile using cross-section analysis. We measured the height values in the x, y directions in the area around Tyr-3NT residues (Fig. 5b, Fig. S5) as well as for the entire surface of the G-actin monomer in the initial conformation, collecting each of the three pixels along x-direction (Fig. 5c). A cross-section analysis around oxidized and non-oxidized Tyr residues in the simulated HS-AFM images of G-actin monomers revealed a difference in the height values from ∼0.2-0.6 nm in magnitude between two G-actin structures, one with non-modified Tyr residues and one with Tyr-3NT residues, respectively (Fig. 5b, lower panel). Rotation of these two structures with 45° step around x-axis or y-axis revealed similar magnitude in differences (Fig. S5).

**Fig. 5.**
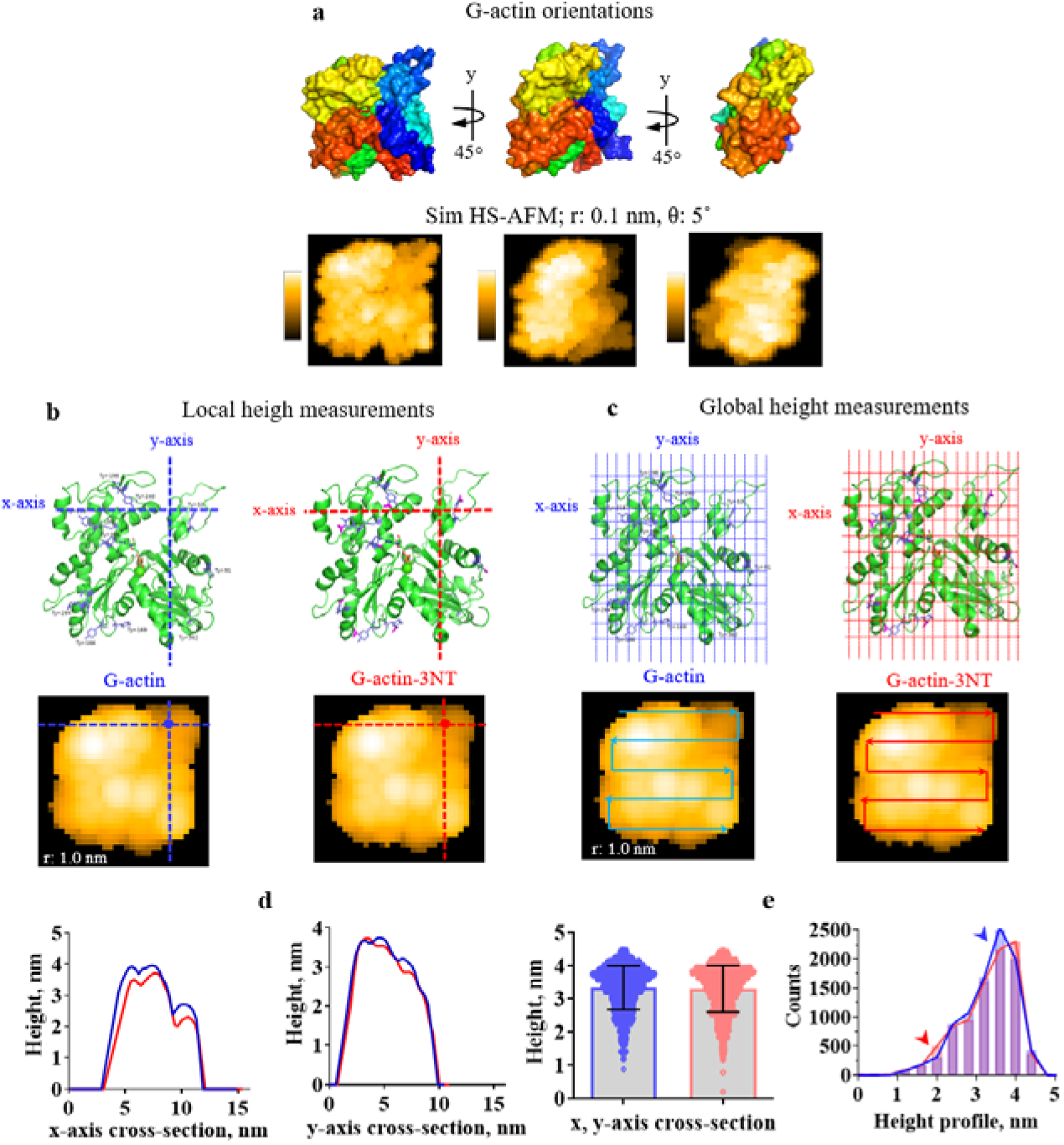
Cross-sectional analysis in the simulated G-actin and G-actin-3NT HS-AFM images. **(a)** The initial conformation of the G-actin monomer structure (PDB: 2ZWH) rotated with 45° step around y-axis and corresponding simulated HS-AFM images (Sim HS-AFM) obtained with BioAFM viewer with the tip radius 0.1 nm and the apex angle 5°. **(b, c)** G-actin structures with non-oxidized Tyr and Tyr-3NT residues, indicating the x, y-axis used to obtain the local height values **(b)** and global height values **(c)** with corresponding simulated HS-AFM images of the G-actin structures with tip radius 1.0 nm with schematic directions of the scanned surfaces. **(d, e)** local and global height profiles for the G-actin structures with non-oxidized Tyr (blue) and Tyr-3NT (red) residues with the frequency analysis of the height values from entire surface. The red arrow shows a higher frequency of 1.8-2.2 nm height values in the oxidized G-actin-3NT; the blue arrow shows the higher frequency of 3-4 nm he ght values in the non-oxidized G-actin.

The entire surface scan of the simulated HS-AFM image of the G-actin monomer in the initial position (Fig. 5e) showed an interesting trend in the height distribution. Despite a similar mean value of 3.3 nm ± 0.66 nm (G-actin) and 3.3 nm ± 0.70 nm (G-actin-3NT), the height frequency comparison revealed that G-actin-3NT had a greater occurrence of low height values (∼1.8-2.2 nm) compared to G-actin, which showed a higher frequency of values ranging from 3.5-4.0 nm (Fig. 5e, the height distribution panel). Thus, according to the computational analysis, the 3-NT oxidative nitration of the Tyr residues contributes to the instability of the system (Fig. 4) that can be evaluated in HS-AFM experiments by the height distributions analysis. Our next goal was to scale the observed differences on the F-actin structure using computational and experimental approaches.

### Global fitting of experimental data for simulation of HS-AFM images

As discussed above, the molecular orientation of G-actin structure for cross-sectional analysis was selected from the molecular dynamics (MD) simulation (Fig. 4). To validate the conformations of the actin filaments in experimental situations, tyrosine residues were modified by 3-NT oxidation and incorporated into the F-actin structure (Fig. 6, Movie S8). This F-actin structure (PDB 6BNO) contains four G-actin monomers for each actin helix, corresponding to ∼22 nm in the length of the actin filament.

**Fig. 6.**
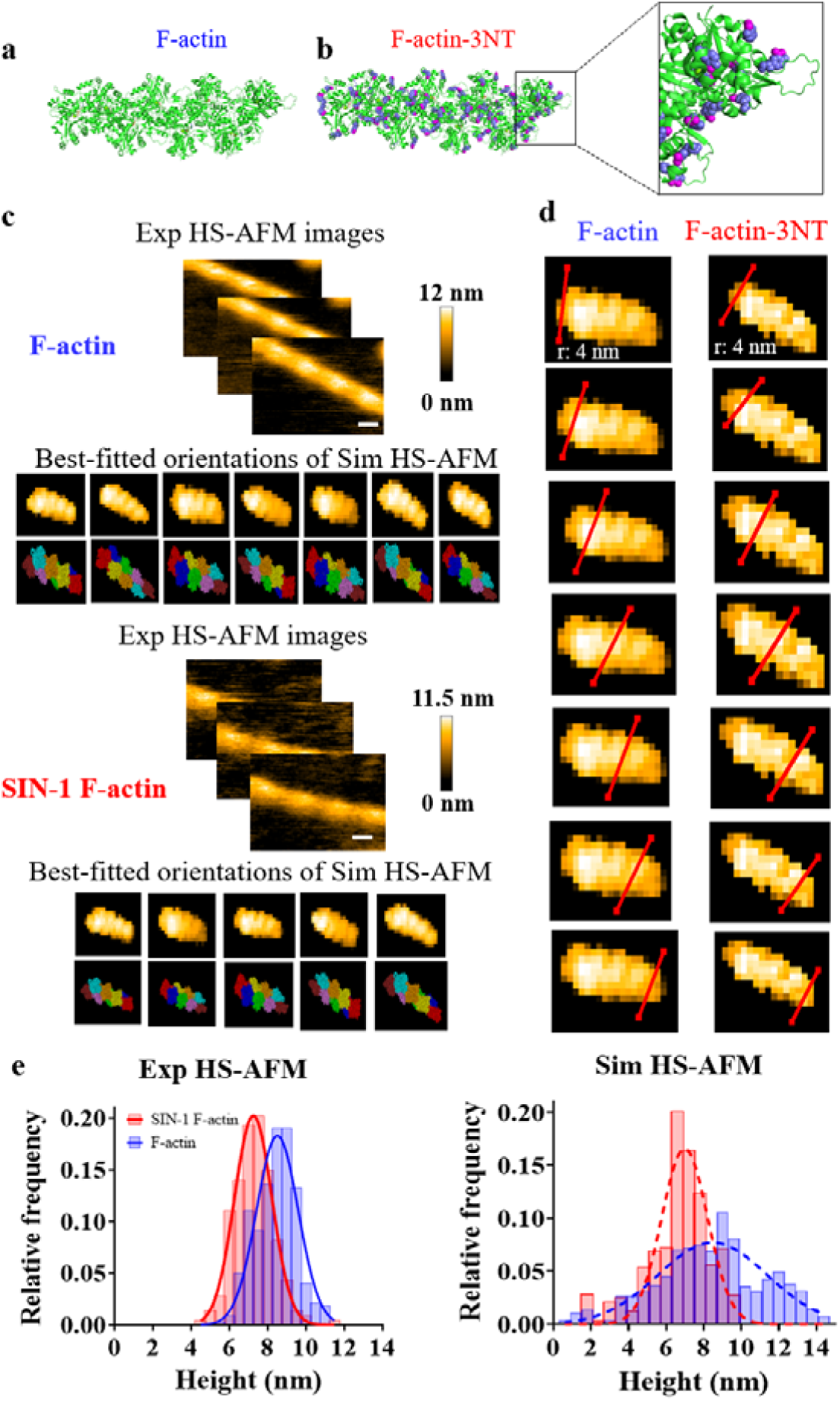
Global fitting of experimental (Exp) and simulated (Sim) HS-AFM images. **(a)** The non-oxidized and **(b)** computationally oxidized F-actin (PDB 6BNO). The modified tyrosine residues are shown as blue spheres. Oxidative modifications were performed in the pyMOL using pyTM plugin (v.1.2)**^30^**. The oxidized atoms in the residues highlighted by pink color. **(c)** Best-fitted orientations of the simulated actin filaments with experimental HS-AFM images of the non-oxidized F-actin and the oxidized by SIN-1 F-actin used as a template. **(d)** An example of the cross-section analysis for the simulated F-actin structure (PDB 6BNO) with non-modified Tyr residues and oxidized Tyr-3NT residues. **(e)** Comparison of the height profiles of the actin filaments in the experimental and simulated HS-AFM images.

The experimental HS-AFM images of the non-treated and SIN-1-treated F-actin were used as a template to gain the best-fitted orientations of simulated F-actin (PDB 6BNO) using global fitting approach available in BioAFM software (see Methods). The resulted best-fitted orientations of the non-oxidized and the oxidized F-actin shown in Figure 6: up to 5-7 different best-fitted orientations were considered to obtain the height profiles of the simulated images. The best-fitted orientations for each experimental HS-AFM datasets were obtained and used for cross-sectional analysis (Fig. 6).

The data for each best-fitted orientation were merged and plotted (Fig. 6e, right). The height profile from simulated HS-AFM images showed very similar height distribution in comparison to the experimental height distributions: 8.5 nm ± 1.1 nm (SD) for non-treated F-actin and 7.2 nm ± 0.97 nm (SD) for SIN-1 oxidized F-actin. The similarity between values measured in the experiments and in the simulated best-fitted orientations validates our computational approach to study conformational features of the biomolecules modified by oxidation conditions.

## DISCUSSION

### SIN-1-induced structural changes in myosin heads

In this study, we employed a combination of computational analysis and high-speed Atomic Force Microscopy (HS-AFM) to investigate the influence of oxidative modifications on the structure-function relationship of the actin-myosin complex. Our study revealed the significant impact of oxidative modifications on the dynamic interplay between myosin and actin at a single-molecule level. We observed alterations in the structural orientation of HMM heads following treatment with SIN-1. According to our HS-AFM data, the oxidized myosin heads exhibited interconnection with each other and potentially with the myosin proximal S2 fragment during the SIN-1 treatment (Fig, 1, Movie S1). This conformational shift could be responsible for the decreased functional activity of oxidized myosin. It was shown that SIN-1 treatment of both myosin and F-actin in the separate experiments, led to a decreased myosin-propelled sliding velocity of actin filaments**^12^**, and reduced force-generation of skeletal myofibrils exposed to SIN-1**^15^**. Moreover, studies demonstrated that oxidation-induced inhibition of muscle fiber contractility is associated with changes in the structural states of myosin**^13,34^**.

Notably, the structural orientation of the oxidized skeletal myosin heads (Fig. 1, Movie S1) displayed a similarity in structural morphology to the super relaxed state (SRX) of cardiac myosin**^35,36^** (Fig. 7). The super relaxed state of myosin is characterized by significantly slow ATP turnover kinetics due to the autoinhibition of the myosin catalytic domains. The autoinhibition of myosin heads in the SRX state is proposed to result from a folded state, where the asymmetric interaction between the two myosin heads and S2 tail is known as the interacting heads motif (IHM)**^36^**.

**Fig. 7.**
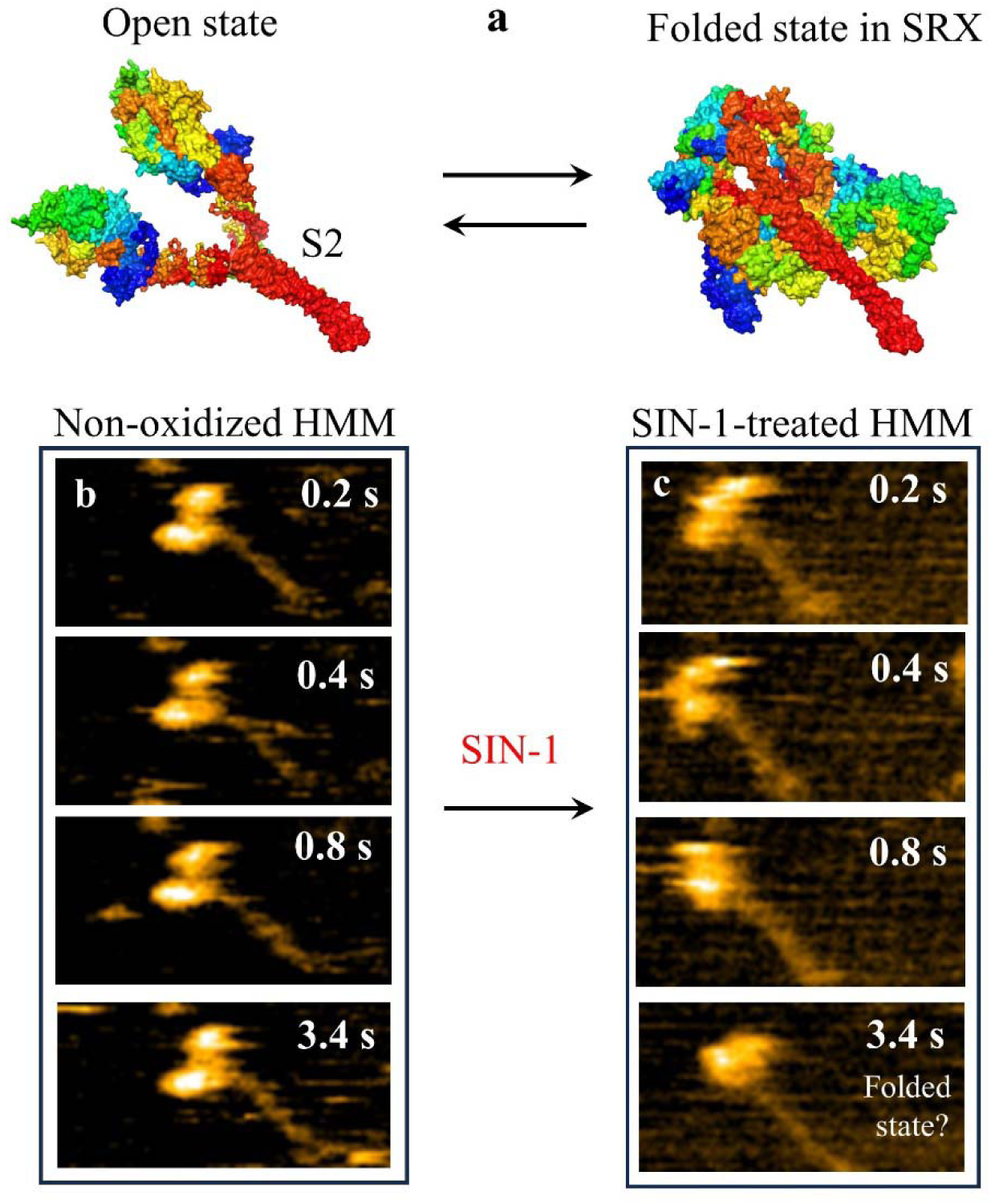
Structural models of the myosin N-terminal domain in the open and closed (super relaxed) states and HS-AFM images of non-oxidized and SIN-1 treated skeletal HMM. **(a)** The non-oxidized model of skeletal myosin (PDB 2MYS, open state) and an energy-conserving, super relaxed state model of human cardiac myosin, MS03 (closed state)**^35^ (b)** The non-oxidized and **(c)** SIN-1-treated skeletal HMM on the mica-APTES surface. Scan area: 150 × 75 nm^2^, 80 × 40 pixels^2^, temporal resolution: 5 frame per second, Scale bar: 30 nm.

Based on HS-AFM images and movies, the treatment of skeletal myosin with SIN-1 has been observed to induce the interaction between the two heads of myosin and possibly with the S2 fragment (Figs 1, 7, Movie S1). Therefore, the proposed similarity in the folded states of myosin in the SRX and under SIN-1 oxidative stress may have significant implications for understanding various pathological oxidative conditions.

### SIN-1-induced structural changes in actin filaments

On the other hand, when SIN-1-treated actin filaments were used to form a complex with non-oxidized HMM, we observed impaired myosin kinetics, established as a reduced size of the HMM head displacement on the actin filament. The displacements of the myosin heads on the F-actin in the presence of ATP suggests that the power stroke is occurring in a series of steps: (i) the myosin molecule binds to the F-actin (M.ADP.Pi); (ii) the myosin lever arm rotates, inducing Pi release (M.ADP); (iii) the myosin molecule detaches from the F-actin (M.ATP), and (iv) the myosin molecule re-binds to the next binding site on the F-actin (M.ADP.Pi). Typically, under our experimental conditions (low ionic strength with 25-100 mM KCl, 2-4 mM MgCL_2_, 1 mM EGTA and neutral pH 7.0), the displacement distribution of the myosin heads bound to the non-oxidized F-actin showed three common sizes: short displacements (under 1 nm), moderate displacements (∼1-3 nm) and long displacements (over 3-4 nm). According to various experimental data, the long displacements ranged from 3-4 nm and over with averaged values ∼4.0-6.1 nm, as in our study, most likely represent the Pi release step. The short displacements ranged from 0.2-2.9 nm with averaged values ∼1.7-2.6 nm most likely represent the ADP release step**^19–22^**. This pattern of displacement distribution was also observed in our previous studies, where non-oxidized actin-myosin complex was used**^16,17^** and showed similar displacement size with experimental data for cardiac and skeletal myosins**^23^**. We found that the displacement distribution of the HMM molecules complexed with SIN-1-treated F-actin revealed the shift to shorter displacements, suggesting that the frequency of long displacement events was significantly decreased by the oxidation (Fig. 1d). According to numerous studies the long displacements related to the phosphate (Pi) release step and lever arm movements responsible for force-generating step**^16,19,23,37^**. Therefore, the observed results suggest that SIN-1 oxidative treatment decreases the pool of the weak bound myosin molecules and shortens the long displacements related to the Pi release step, thus reducing the force generation by myosin motors.

The reduced size of HMM heads displacement in the SIN-1-treated F-actin-HMM complex can be explained by oxidation-induced conformational changes in actin filaments. For instance, several studies have demonstrated that the treatment of F-actin with SIN-1 leads to the inhibition of actin polymerization. This inhibition has been shown to be concentration-dependent**^2,13^**, resulting in a significant reduction of the ability to stimulate actin-activated myosin ATPase activity**^13^**. We approached to compare the non-oxidized and SIN-1-oxidized actin structures using HS-AFM that included the assessment of actin peak heights and the half-helical pitch, i.e. the distance between each peak. Previously, observation of the actin topology under the HS-AFM has revealed changes at actin structure in response to binding partners such as Ca^2+^ ions binding to the thin filaments**^37^**, cofilin binding to the actin filaments**^38^**, myosin binding to actin filaments**^26,17^** or actin-binding domain of Rng2**^39^**. The peaks in the F-actin structure captured by HS-AFM represent the highest points of the actin helix, where two strands of the actin filament are aligned atop each other. Due to the large size and highly dynamic nature of actin filaments, achieving high resolution in the area, where two stands are crossed is challenging (Fig. 2b). Therefore, the highest point represents the area of actin structure that includes at least 14 G-actin monomers (7 monomers for each actin strand). Therefore, the treatment of actin filaments with SIN-1 leads to a reduction in the peak height distribution in comparison to non-oxidized filaments (Fig. 2d), suggesting oxidation-induced conformational changes in the orientation of the G-actin monomers.

Importantly to note that SIN-1 treatment specifically increases the amount of nitrated tyrosine residues (3-NT), caused by the addition of a nitro (-NO2) group in ortho position of the aromatic cycle, while not impacting other redox post-translational modifications (PTMs) such as dityrosine or carbonyl formation**^40,41^**. Additionally, 3-NT tyrosine nitration (NO2; + 46 Da) is irreversible modification**^40^** that has been found to be exclusively associated with SIN-1 treatment. 3-NT modifications were consistently observed in myofibril actin and purified skeletal muscle F-actin obtained from both rheumatoid arthritis (RA) patients and mice with experimentally induced RA models**^2^**. The presence of 3-NT and MDA modifications in proteins oxidized by SIN-1 is believed to hinder intramolecular hydrogen bonding, potentially leading to conformational changes within the actin structure**^2,29,40–42^**.

### Computational analysis of 3-NT modifications on the G-actin and F-actin structures

To gain further insights into the structural changes induced by SIN-1 and specifically by tyrosine nitration in actin, we employed molecular dynamics (MD) and HS-AFM simulations, establishing a connection to HS-AFM experiments on SIN-1-oxidized F-actin structure. Despite the fact that only one type of oxidized residue, tyrosine, is present in our computational model, does not weaken the model and comparison with experimental data. Instead, it enables us to examine in detail the impact of tyrosine nitration that is linked to a wide range of disorders. Tyrosine nitration is a post-translational modification that contributes to the development of protein disfunction**^42^**, having an impact on muscle contractile function as demonstrated with *in vitro* studies**^43–45^** and various pathological conditions, including neurodegenerative, cardiovascular and autoimmune diseases, such as rheumatoid arthritis**^2,43^**. We observed that targeted tyrosine residues in G-actin structure undergo oxidative modification destabilized the protein system (Fig. 4). With AI-Copilot model (www.310.ai) we showed that several tyrosine residues are involved in the binding pockets of the actin-tropomyosin-myosin-MgADP complex (PDB: 8EFH) (Supplementary Table 1, Fig.S6). Simulated HS-AFM images of G-actin and F-actin were analyzed to test structural modifications that occurred due to the nitration of tyrosine residues, by examining the peak heights distribution in the non-oxidized and oxidized proteins. The decrease in height distribution in the computationally oxidized G-actin / F-actin structures simulated as HS-AFM images was in the same range as was observed in our experimental HS-AFM data, indicating a consistent observation of the difference in height distribution.

### Summary

In this paper we demonstrated that oxidative modifications induce structural changes in the skeletal muscle myosin heads and actin filaments. Specifically, the oxidized HMM molecule exhibited a significant alteration in the spatial orientation of the myosin heads compared to the non-oxidized molecules. This change in head orientation affected the binding capacity of the oxidized HMM to non-oxidized actin filaments in HS-AFM experiments. Conversely, when oxidized actin filaments were combined with non-oxidized myosin in the presence of ATP, it resulted in a reduction in the size of myosin head displacement and force generation. Furthermore, we utilized molecular dynamics (MD) simulation and simulated HS-AFM images to investigate the structural impact of oxidative tyrosine nitration on actin subunits, aiming to reveal the functional consequences of this modification on actin’s structure.

## Methods

### Proteins

Skeletal muscle HMM and F-actin were purified and tested for their functional activity by in-vitro motility assays, as described**^37^**. In some experiments skeletal HMM from Cytoskeleton Inc. (USA) was used.

### 3NT-oxidation of the tyrosine residues

The G-actin (PDB 2ZWH) and F-actin (PDB 6BNO) were used as non oxidized molecular structures. The computational oxidative nitration (3NT) of the Tyr residues was implemented in a PyMOL software (v.2.3.3, Schrodinger LLC), using PyTMs plugin**^30^**. The selection of the Tyr residues was based on the mass-spectrometry analysis of F-actin from patients with rheumatoid arthritis in our previous study**^2^**. The analysis showed the hotspots of oxidized residues in each of the subdomain of G-actin.

### Molecular dynamics simulations

The initial atomic coordinates were obtained from the 3.3 Å X-ray structure for skeletal G-actin (PDB 2ZWH). The atomic spatial coordinates of the non-modified G-actin and G-actin-3NT with oxidized Tyr residues were used to obtain trajectory from Molecular Dynamics (MD) simulations to evaluate the electrostatic potential energy and root-mean-square deviation (RMSD). MD simulations were carried out by our custom Python code (GitHub: <u>matusoff/Molecular-dynamics-with-BioPython)</u> using OpenMM package (http://docs.openmm.org/latest/api-python/)**^31^** installed in the conda environment in Python. The TIP3P water model (’tip3p.xml’) and the AMBER forcefield (’amber99sbildn.xml’) from OpenMM module were used.

The non-oxidized G-actin and oxidized G-actin-3NT structures contain 5894 atoms and 5960 atoms, respectively. To pre-process the PDB files before MD simulations (add missing residues, add missing atoms, remove hydrogens, fix positions, get topology) we used PDBFixer.py script, based on OpenMM package, which is available on the GitHub: (https://github.com/matusoff/Molecular-dynamics-with-BioPython/blob/main/PDBFixer.py). The non-oxidized G-actin or oxidized G-actin-3NT structures were placed in a rectangular box, filled with water molecules using VMD software (v.1.9.3)**^46^**. The size of the box defined by VMD software based on molecular dimensions of the protein structure: G-actin: 58.4 Å × 64.0 Å × 71.5 Å, G-actin-3NT: 83.2 Å × 88.8 Å × 96.5 Å. The protein atoms to its boundaries were at least 1.5 nm, and periodic conditions were set at the borders. Na+ and Cl− ions were added to ensure the electrical neutrality of the system and the ionic strength was settled up to 0.15 M (Movie S6). The charge of G-actin and G-actin-3NT before ionization was negative (−12.99); after ionization the charge was neutral (∼4.3-4.5e^-6^).

Energy minimization was carried out using the OpenMM module using force groups defined by the LangevinIntegrator class. The process included finding a local minimum of the potential energy of a molecular system to optimize the initial structure. After energy minimization, the system was equilibrated in the NVT (constant number of particles, volume, and temperature) and NPT (constant number of particles, pressure, and temperature) ensembles. The system is balanced to a temperature of 300 K. The duration of the MD trajectories was settled up to 100 ns with a step of 1 ps. Following constants for the MD simulations were used: Boltzmann constant in kcal/mol/K: k = 1.987e-3; temperature in Kelvin: 300.0, number of MD steps to run:100000. The root-mean-square deviation (RMSD) was calculated in the VMD software with obtained during MD simulation trajectory dcd file.

### Simulation of the HS-AFM images

Simulation of the HS-AFM images of computationally oxidized and non-oxidized G-actin and F-actin structures was done in BioAFM viewer (v.2.5, Kanazawa University). The following protocol was used: (i) Load PDB file with the non-modified Tyr residues or PDB file with the Tyr residues modified by oxidative nitration. The initial position of the atomic structure for G-actin was as shown in Fig. 1, where all four G-actin subdomains are visible. To ensure similar orientation of the two loaded structures, the correlation error was calculated by the custom Python script available in Supplementary Data; (ii) Rotation of the structure from the initial position by the step of 45° degree around y-axis or around x-axis; (iii) Cross-sectional analysis of the area around non-modified residues and residues modified by oxidative nitration (horizontal and vertical cross-sections).

As soon the initial position is settled up, we save the initial position coordinates for further use. To obtain different structural orientation the rotation around x or y-axis was performed at 45° degree step. The structural orientations shown in Figures 3, S1 and Movie S2. To compare the simulated non-oxidized and oxidized HS-AFM images correctly, the rotation of each image must be done to the exact degree step from the initial position. The spots around the non-oxidized and the oxidized Tyr residues were analyzed in the simulated HS-AFM images for the peak height distributions and cross-section analysis.

### SIN-1 treatment of the skeletal F-actin and myosin HMM molecules

The SIN-1 treatment of F-actin and myosin HMM was performed according to our previously published paper**^12^**. Briefly, freshly prepared SIN-1 was incubated at room temperature for 15 min and added to 7 μM F-actin in a buffer 60 mM KCl, 0.1 mM EGTA, 3 mM NaN_3_, 2 mM MgCl_2_, 10 mM MOPS at pH 7.0. and allowed to react for 10 min. The SIN-1-treated F-actin was stored at −80 C before use in the HS-AFM experiments. The HMM molecules were treated by SIN-1 in the following conditions: freshly prepared SIN-1 was incubated at room temperature for 15 min and added to 4 nM HMM (Cytoskeleton, USA). The final concentration of SIN-1 was 60 μM.

### Myosin displacement analysis and force calculation

The definition of the power stroke during the HS-AFM acquisition of the actin-myosin complex and the approach to measure the myosin displacements described in detail in our previous works**^16,17,47^**. Here, we compared the myosin HMM displacements along the non-oxidize and the oxidized actin filaments attached to the mica-SLB surface in the presence of 2 µM ATP. The myosin HMM displacement was calculated as the distance between the averaged center of mass (COM) position of HMM heads bound to F-actin in the reference frame and in the next frames, in successive HS-AFM images. The COM of the HMM head was calculated by the x, y, and z coordinates of the HS-AFM images. The x and y data correspond to the lateral coordinates, while the z coordinates described the height of the highest point in the HMM heads. The polarity of actin filament defined by the orientation of bound HMM head as described**^47^**.

The force estimation of the HMM heads bound to non-treated or SIN-1-treated F-actin was done with following equation:

*F = k \** Δ*d*, where d is the displacement of HMM heads on actin filaments measured in the present study and *k* is a stiffness of myosin molecule (2 pN/nm), calculated based on experimental data and computational modelling of the free energy profiles of the attached cross-bridges (see **^18^**).

### HS-AFM and cantilevers

The experiments were performed on a tapping-mode HS-AFM system (RIBM)**^48^** by using small Olympus BL-AC10DS-A2 cantilevers with the following parameters: spring constant: 0.08-0.15 N/m, quality factor in water: ∼1.4-1.6, resonant frequency in water: 06-1.2 MHz. The probe tip was fabricated on the tip of a cantilever by electron-beam deposition and sharped by plasma etcher with a ∼4 nm tip apex. A tip–sample loading force can be modulated by the free oscillation peak-to-peak amplitude (A_0_) of the cantilever set to ∼2.0 nm and the amplitude set point adjusted to more than 0.9 A_0_.

### HS-AFM sample preparation

The lipid composition used for HS-AFM imaging contained 1,2-Dipalmitoyl-*sn*-glycero-3-phosphocholine (DPPC, Avanti Polar Lipids), 1,2-Dipalmitoyl-3-trimethylammonium-propane (DPTAP, Avanti Polar Lipids) and 1,2-dipalmitoyl-*sn*-glycero-3-phosphoethanolamine-*N*-(cap biotinyl) (biotin-cap-DPPE, Avanti Polar Lipids). DPPC: DPTAP: biotin-cap-DPPE were mixed in a weight ratio of 89:10:1. The preparation of lipid bilayer vesicles and deposition on mica substrate has been previously described**^16^**.

### HS-AFM imaging

In order to assess possible structural changes in F-actin after exposure to SIN-1 modification, HS-AFM experiments were conducted with immobilized F-actin on a mica-supported lipid bilayer substrate (mica-SLB), containing a mixture of negatively (DPPC) and positively (DPTAP) charged lipids with small fraction of biotin-cap DPPE (ratio: 89:10:1). This lipid substrate allowed actin to attach to the surface, while maintaining some freedom of movement**^16,37,49^** which was essential for the analysis of actin structure and oxidation-induced structural changes.

After rinsing the substrate with attachment buffer (25 mM KCl, 2 mM MgCl_2_, 1 mM EGTA, 25 mM Imidazole-HCl, pH 7.0), 3.0 µl of 7 µM non-treated or SIN-1-treated actin filaments diluted in attachment buffer was deposited on the mica-SLB substrate and incubated for 10 minutes in a wet chamber. The unbounded actin filaments were washed by the same solution, and 3 µl of 8 nM HMM diluted in attachment buffer was placed on top of the bound to the substrate actin filaments and incubated for an additional 5 minutes. The actin-HMM complex in nucleotide-free conditions was rinsed by 20 μl of attachment buffer, containing 2 µM ATP dissolved in attachment buffer.

The sample stage with the mica-SLB substrate, containing actin-HMM complex was immersed into the AFM liquid cell chamber (volume ∼120 µl) filled with the solutions matching the experimental conditions. HS-AFM observations were performed at 5-10 frame per second (fps) with a typical scan area of 150 nm in x-direction and pixel resolutions of 80 x 40 pixels^2^. The specific pixel resolutions and scan area range are indicated in the figure and movies legends. All experiments were performed at room temperature (25°C).

### Data analysis and processing of HS-AFM images

The HS-AFM images were analyzed by Kodec software (4.4.7.39)**^38^** and imageJ, with a low-pass filtering to remove spike noise in the image and to make the *xy*-plane flat. To simplify the analysis, we chose actin filaments located approximately in parallel to the x-direction. HMM heads bound to actin were visualized as a globular sphere of ∼20 nm in diameter with the neck part, containing the ∼8-12 nm lever-arm region. Statistical analysis (student t-test, correlation), Gaussian distributions and fitting were performed in GraphPad Prism software (v.9.3.0). Values are reported as mean ± Standard Deviation. A level of significance of P ≤ 0.05 was set for all analyses.

## Data availability

All data required for evaluation of the conclusions in the paper are present in the main body of the paper and/or in the Supporting Information. Additional data are available related to this paper may be requested from the authors.

## Acknowledgments

This work was supported by the Natural Science and Engineering Research Council of Canada (to D.E.R). D.E.R. is a Canada Research Chair in Muscle Biophysics.

## Author contributions

O.S.M. and D.E.R. designed research; O.S.M. performed HS-AFM experiments and created custom Python codes to pre-process files before MD simulations and Python codes to perform MD simulations using OpenMM environment. All authors were involved in analysis and interpretation of the data. O.S.M., D.E.R. wrote the paper and all authors approved the final version of the manuscript.

## Ethics declarations

Competing interests

The authors declare no competing interests.

## Notes

### Competing Interest Statement

The authors have declared no competing interest.

