## Supplementary Figures for "Oxidation-induced structural changes in actin and myosin evaluated by computational simulation, machine learning modeling and high-speed AFM"

**Supplementary file includes:**

Python code for Molecular dynamics simulation

Python scripts for image analysis

Table S1

Figures S1 to S6.

Captions for Movies S1 to S9.

**Other Supplementary Materials for this manuscript include the following:**

Movies S1 to S9

The custom Python code used for Molecular dynamics (MD) simulation with OpenMM package and Amber forcefield.

**OpenMM_Simulation.py**

import numpy as np

import matplotlib.pyplot as plt

import mdtraj as md

import Bio.PDB

from Bio.PDB import PDBParser

import simtk.unit as unit

import openmm as mm

import openmm.app as app

from openmm.app import *

from openmm import *

from simtk import unit

from sys import stdout

import nglview as nv

\# Load the PDB file

pdb = PDBFile(r'path\2zwh_fixed_pH_7.pdb')

k = 1.987e-3 # Boltzmann constant in kcal/mol/K

T = 300.0 # temperature in K

dt = 0.002 # integration timestep in ps

nsteps = 100000 # number of MD steps to run

\# Define the force field

forcefield = app.ForceField('amber99sbildn.xml', 'tip3p.xml')

\# Create the simulation system

system = forcefield.createSystem(pdb.topology, nonbondedMethod=app.PME, nonbondedCutoff=1.2*unit.nanometers, constraints=app.HBonds)

box_vectors = np.diag([10, 10, 50]) * unit.nanometer

system.setDefaultPeriodicBoxVectors(*box_vectors)

\# Add a Langevin thermostat

integrator = LangevinIntegrator(T*unit.kelvin, 1.0/unit.picosecond, dt*unit.picoseconds)

integrator.setRandomNumberSeed(42)

\# Create the simulation object

platform = mm.Platform.getPlatformByName('CPU')

simulation = Simulation(pdb.topology, system, integrator, platform)

\# Set the initial positions of the atoms

positions = pdb.getPositions()

simulation.context.setPositions(positions)

\# Minimize the energy

print('Minimizing energy...')

#simulation.minimizeEnergy()

simulation.minimizeEnergy(tolerance=1.0*unit.kilojoule_per_mole, maxIterations=100) #change it to 100 from 1000

\# Equilibrate the system

print('Equilibrating...')

simulation.context.setVelocitiesToTemperature(T*unit.kelvin)

simulation.step(100000)

#Create the DCD reporter to save the trajectory data

report_interval = 1

reporter = DCDReporter(r'path\trajectory.dcd', report_interval)

simulation.reporters.append(reporter)

\# Run the production simulation and calculate the total potential energy at each frame

print('Running production simulation...')

potential_energy = []

for i in range(nsteps):

simulation.step(1)

state = simulation.context.getState(getEnergy=True)

potential_energy.append(state.getPotentialEnergy().value_in_unit(unit.kilocalorie_per_mole))

\# Save the force data to a csv file

np.savetxt('potential_energy.csv', potential_energy, delimiter=',')

\# Plot the potential energy data

plt.plot(range(nsteps), potential_energy)

plt.xlabel('Time (ps)')

plt.ylabel('Potential energy (kcal/mol)')

plt.show()

**PDF_Fixer.py**

This script was used to correct PDB files of G-actin before MD simulation.

from pdbfixer import PDBFixer

from simtk.openmm.app import *

from simtk.openmm import *

from simtk.unit import *

import os

def fix_pdb(pdb_id):

pdb = PDBFile(pdb_id)

if len(pdb_id) != 4:

print("Creating PDBFixer...")

fixer = PDBFixer(pdb_id)

print("Finding missing residues...")

fixer.findMissingResidues()

chains = list(fixer.topology.chains())

keys = fixer.missingResidues.keys()

for key in list(keys):

chain = chains[key[0]]

if key[1] == 0 or key[1] == len(list(chain.residues())):

print("ok")

del fixer.missingResidues[key]

print("Finding nonstandard residues...")

fixer.findNonstandardResidues()

print("Replacing nonstandard residues...")

fixer.replaceNonstandardResidues()

print("Removing heterogens...")

fixer.removeHeterogens(keepWater=True)

print("Finding missing atoms...")

fixer.findMissingAtoms()

print("Adding missing atoms...")

fixer.addMissingAtoms()

print("Adding missing hydrogens...")

fixer.addMissingHydrogens(7)

print("Writing PDB file...")

PDBFile.writeFile(

fixer.topology,

fixer.positions,

open(os.path.join(".", "%s_fixed_pH_%s.pdb" % (pdb_id.split('.')[0], 7)),

"w"),

keepIds=True)

return "%s_fixed_pH_%s.pdb" % (pdb_id.split('.')[0], 7)

**Python code for correlation of G-actin (PDB 2ZWH) and simulated HS-AFM G-actin structure.**

**Script to calculate the mean squared error using neural model to correlate a simulated HS-AFM image and a structure obtained from PDB file**

import numpy as np

import keras

from keras.models import Sequential

from keras.layers import Dense, Conv2D, Flatten

import matplotlib.pyplot as plt

\# Load and preprocess the two images data

\# tip radius for sim AFM = 2.0

\# PDB obtained by Bio-AFM viewer

image1 = np.load('/content/image.npy')

image2 = np.load('/content/image_2zwh-VdW_pdb.npy')

image1 = image1 / 255.0 # normalize to [0, 1] range

image2 = image2 / 255.0 # normalize to [0, 1] range

\# Reshape the images to 4-dimensional tensors

image1 = np.array(image1)

image1 = image1.reshape(100, 100, 1)

image2 = np.array(image2)

image2 = image2.reshape(100, 100, 1)

\# Define the neural network model

model = Sequential()

model.add(Conv2D(32, kernel_size=(3,3), activation='relu', input_shape=(100,100,1)))

model.add(Flatten())

model.add(Dense(512, activation='relu'))

model.add(Dense(256, activation='relu'))

model.add(Dense(128, activation='relu'))

model.add(Dense(64, activation='relu'))

model.add(Dense(1, activation='linear'))

\# Compile the model

model.compile(loss='mean_squared_error', optimizer='adam')

\# Train the model

model.fit(np.array([image1, image2]), np.array([0, 1]), epochs=10, batch_size=32, validation_split=0.2)

\# Use the model to predict the correlation between the two images

correlation_result = model.predict(np.array([image1, image2]))

\# Plot the original images

plt.figure(figsize=(10,5))

plt.subplot(1,2,1)

plt.imshow(image1.reshape(100, 100), cmap='gray')

plt.title("Image 1")

plt.subplot(1,2,2)

plt.imshow(image2.reshape(100, 100), cmap='gray')

plt.title("Image 2")

\# Print the correlation result

print("Correlation result: ", correlation_result)

\# # Normalize the images to [0, 1] range

\# image1 = (image1 - np.min(image1)) / (np.max(image1) - np.min(image1))

\# image2 = (image2 - np.min(image2)) / (np.max(image2) - np.min(image2))

\# Predict the correlation between the two images

prediction = model.predict(np.array([image1, image2]))

\# Merge the two images based on the predicted correlation

merged_image = image1 * prediction[0] + image2 * prediction[1]

\# Plot the merged image

plt.imshow(merged_image.reshape(100, 100), cmap='plasma')

def mean_squared_error(image1, image2):

return np.mean((image1 - image2)**2)

mse = mean_squared_error(image1, image2)

print("Mean Squared Error:", mse)

\# Show the plot

plt.show()

**Results for correlation for G-actin (PDB 2ZWH) and simulated HS-AFM G-actin structure**

Epoch 1/10

1/1 [==============================] - 2s 2s/step - loss: 2.8836e-11 - val_loss: 0.9950

Epoch 2/10

1/1 [==============================] - 1s 762ms/step - loss: 7.2021e-06 - val_loss: 1.0028

Epoch 3/10

1/1 [==============================] - 1s 786ms/step - loss: 2.0976e-06 - val_loss: 0.9979

Epoch 4/10

1/1 [==============================] - 1s 768ms/step - loss: 1.1370e-06 - val_loss: 1.0043

Epoch 5/10

1/1 [==============================] - 1s 771ms/step - loss: 4.7226e-06 - val_loss: 0.9984

Epoch 6/10

1/1 [==============================] - 1s 769ms/step - loss: 6.4588e-07 - val_loss: 1.0097

Epoch 7/10

1/1 [==============================] - 1s 756ms/step - loss: 2.4389e-05 - val_loss: 0.9929

Epoch 8/10

1/1 [==============================] - 1s 763ms/step - loss: 1.2838e-05 - val_loss: 1.0021

Epoch 9/10

1/1 [==============================] - 1s 764ms/step - loss: 1.1487e-06 - val_loss: 1.0008

Epoch 10/10

1/1 [==============================] - 1s 778ms/step - loss: 1.7236e-07 - val_loss: 0.9975

1/1 [==============================] - 0s 175ms/step

Correlation result: [[0.00126135]

[0.00127196]]

1/1 [==============================] - 0s 128ms/step

Mean Squared Error: 6.520016537572202e-07

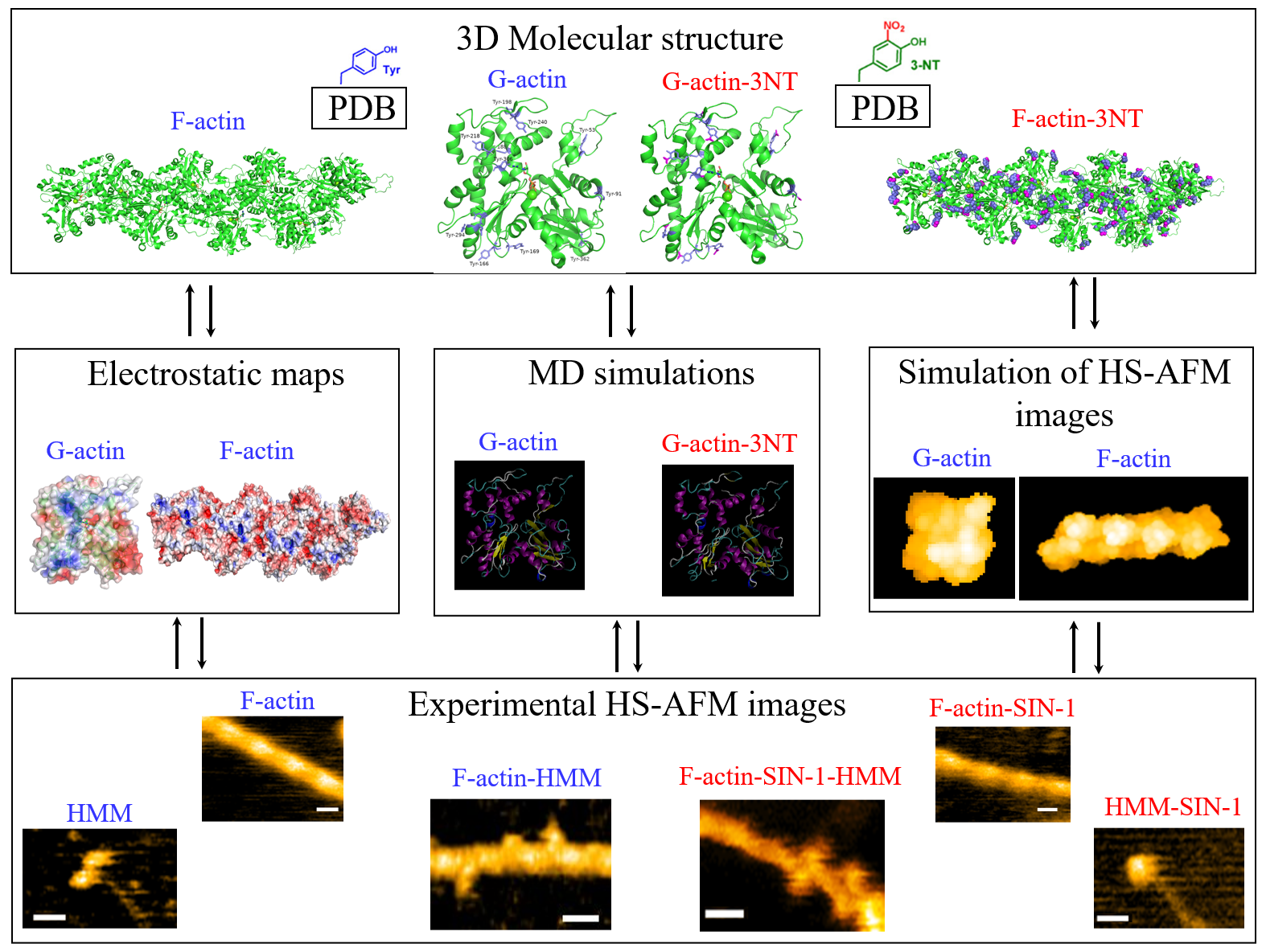

**Figure S1.** **Flowchart of the project.**

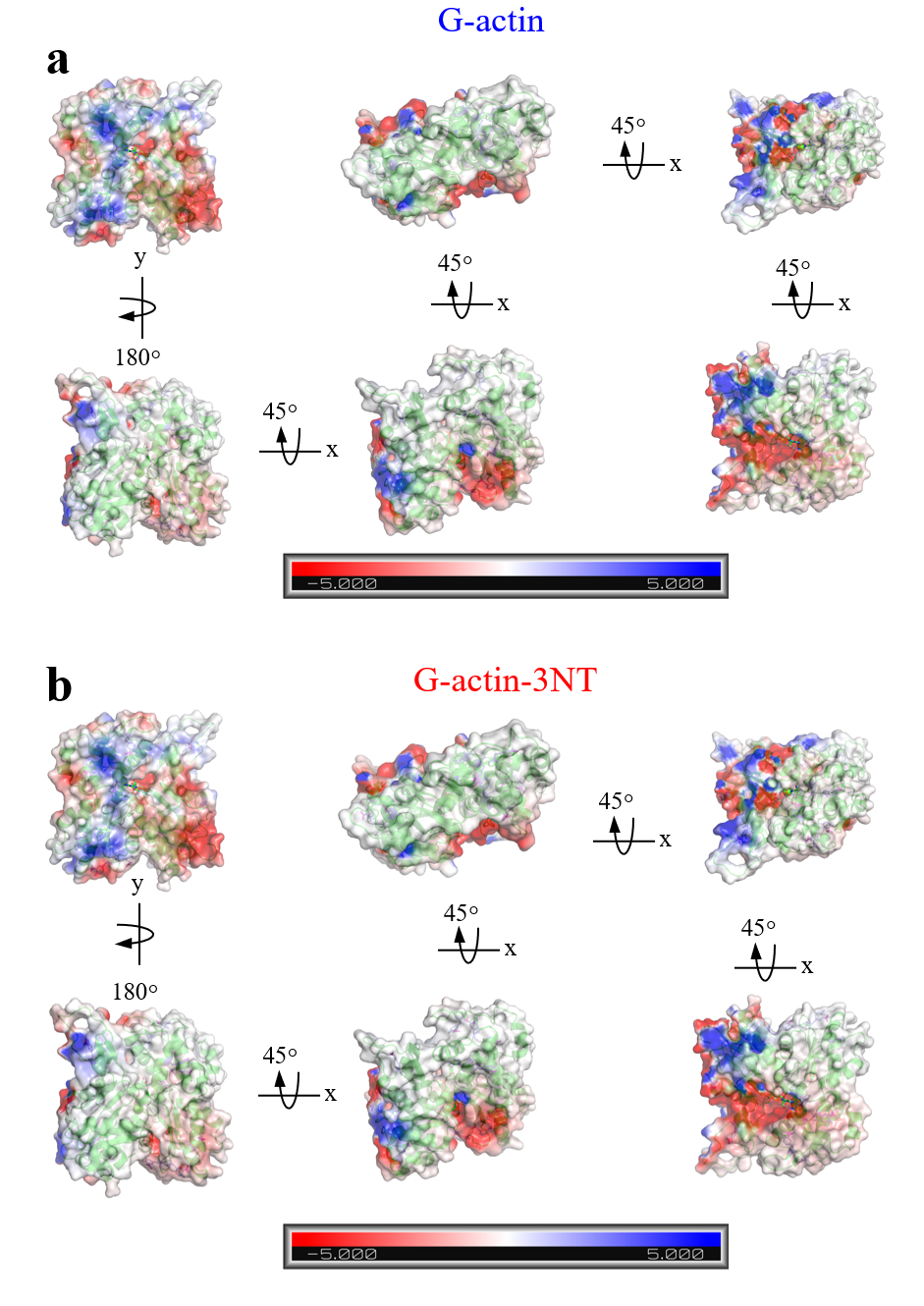

**Figure S2. The successive x-axis rotation of the** **G-actin structures**. The non-oxidized G-actin **(a)** and the oxidized G-actin-3NT **(b)** structures with 45^o^ step from initial position showed the local changes in the electrostatic potential energy around oxidized residues. The matched colored boxes for each orientation were magnified to show the charge differences more precisely. The map scale represents the electrostatic potential energy from -5.0 kBT (red) to 5.0 kBT (blue).

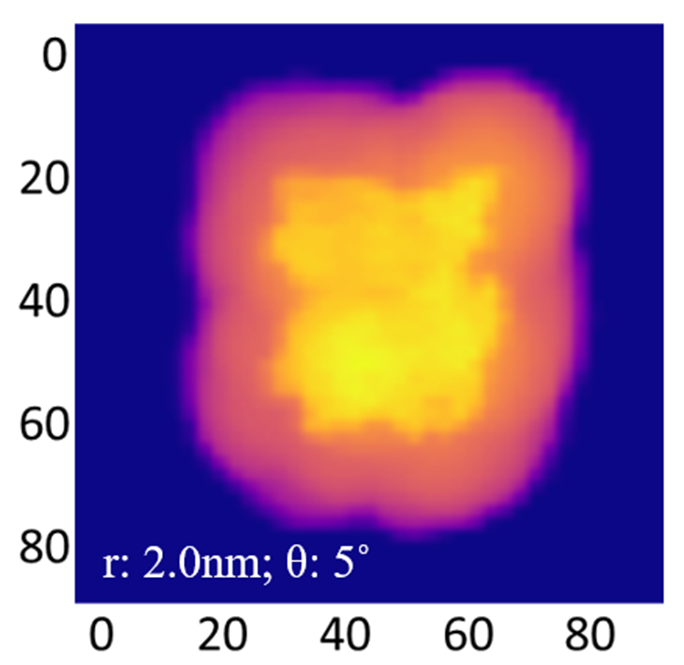

**Figure S3.** **Correlation between pseudo-AFM G-actin image and G-actin structure** **using** **TensorFlow platform**. The pseudo-AFM images were obtained by BioAFM software (v.2.5) as described in the Methods using tip radii 2 nm. The mean squared error: 6.52 _˟_ e-07 indicates the high correlation between pseudo-AFM image and G-actin atomic structure.

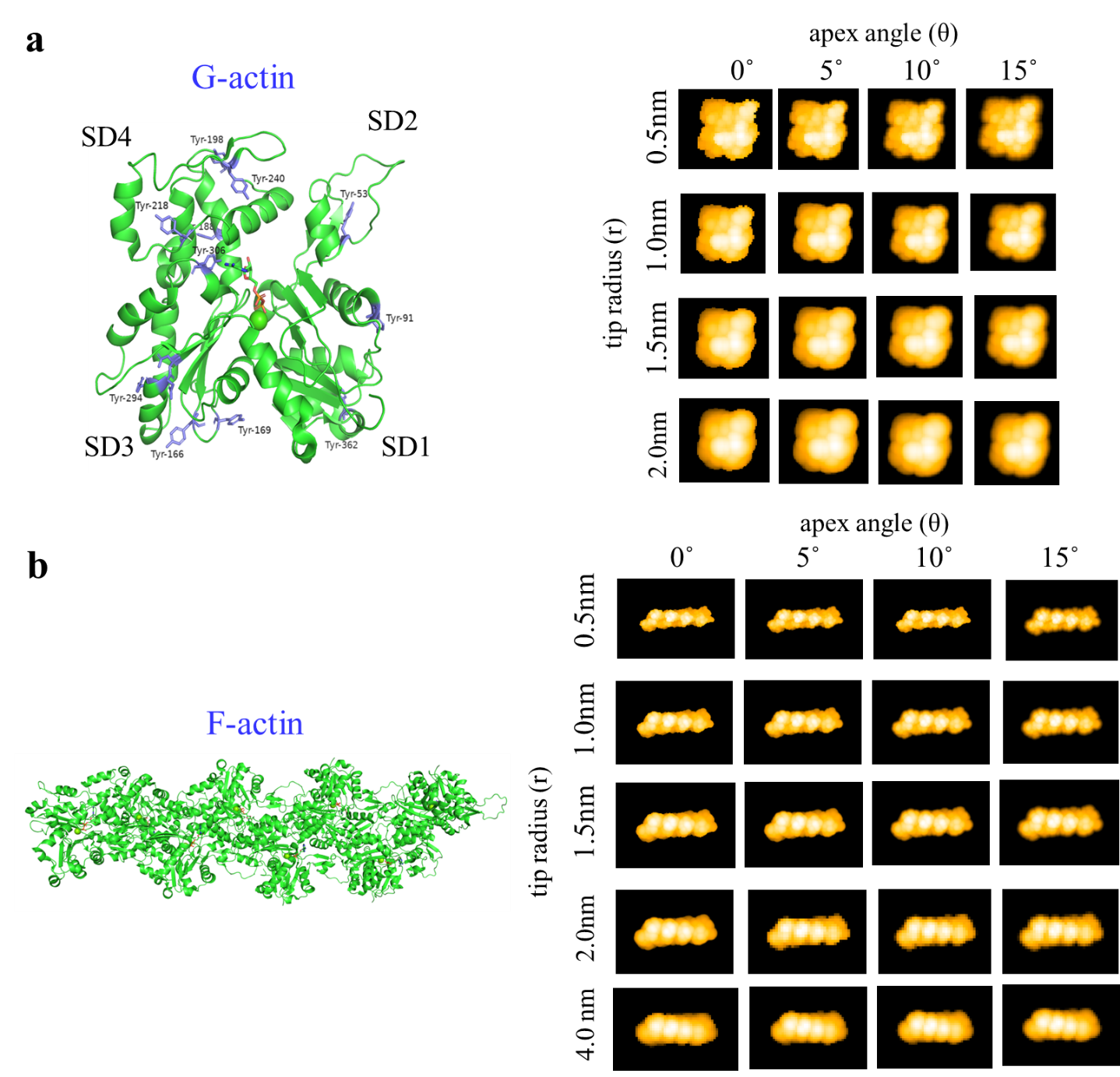

**Figure S4**. **Simulated HS-AFM images captured with different tip radii and apex angles**. The images were captured at tip radii ranging from 0.5 to 4 nm and different apex angles from 0 ^o^ to 15 ^o^, showcasing how the tip makes a contact with the surface.

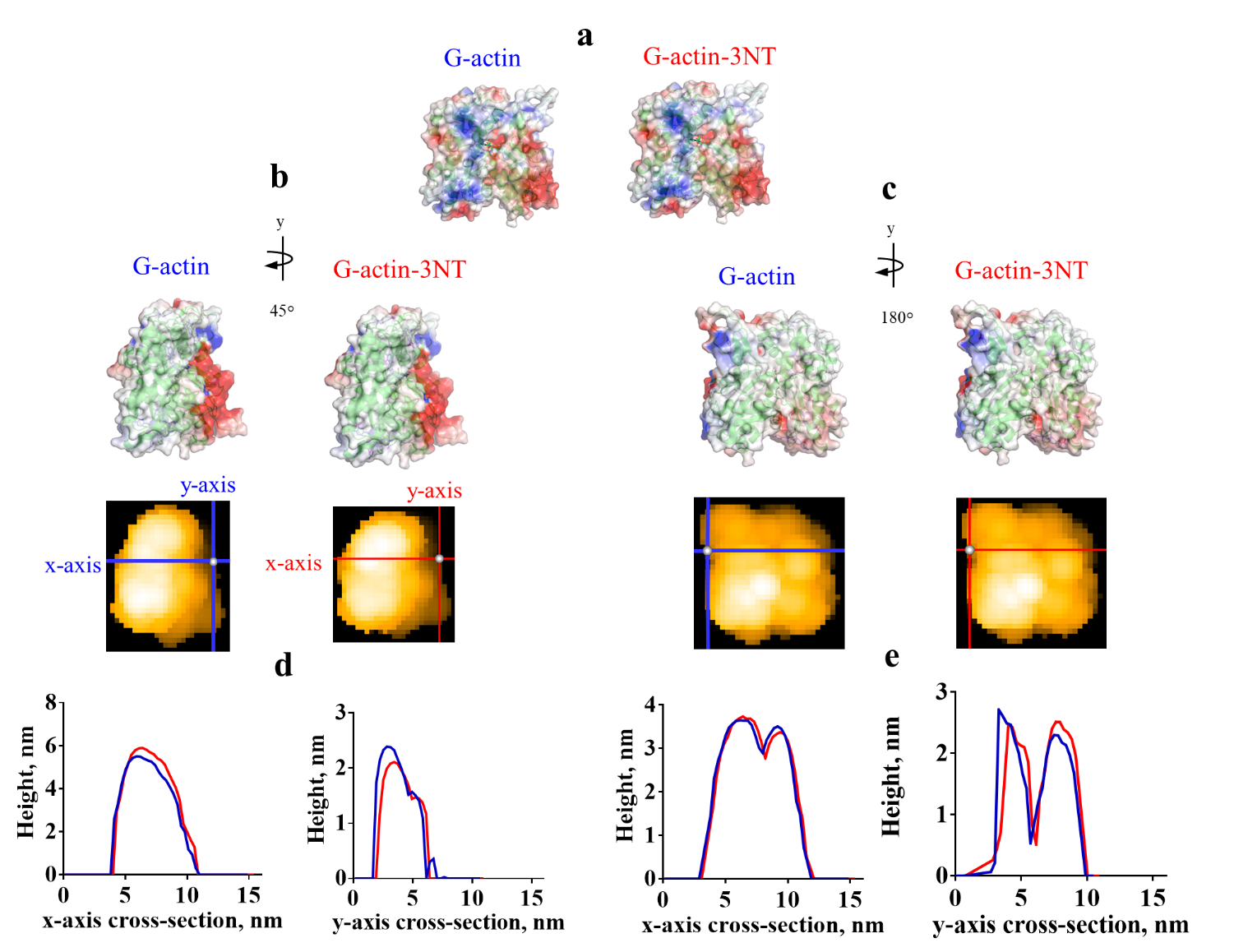

**Figure S5**. **Local height measurements by cross-sectional analysis in the simulated G-actin and G-actin-3NT HS-AFM images. (a-c)** The initial conformation of the G-actin monomer structure (PDB: 2ZWH) rotated with 45^o^ step around y-axis and corresponding simulated HS-AFM images obtained with BioAFM viewer with the tip radius 2 nm and the apex angle 5^o^. G-actin structures with non-oxidized Tyr and Tyr-3NT residues, indicating the x, y-axis used to obtain the local height values. **(d, e)** local height profiles for the G-actin structures with non-oxidized Tyr (blue) and Tyr-3NT (red) residues.

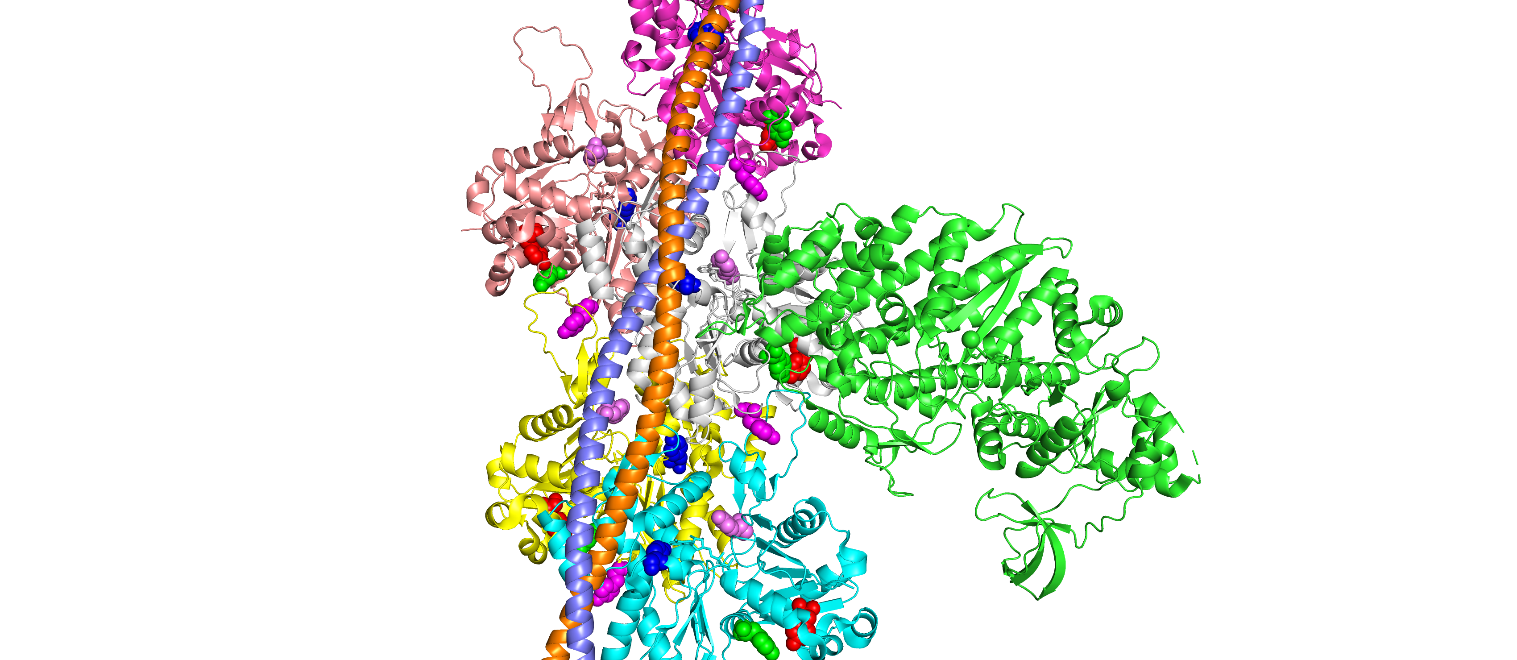

**Figure S6**. **The top five Tyrosine residues located in the binding pockets of actin-tropomyosin-myosin-MgADP complex, 3.3 Å (PDB: 8EFH)^1^ as defined by 310 AI-Copilot model.** The top five Tyrosine residues Tyr133 (red), Tyr306 (blue), Tyr143 (green), Tyr169 (magenta) and Tyr69 (violet) showed the highest probability of binding and indicated as spheres of different colors.

**Supplementary Table 1**

**Table S1. The binding probability scores for each residue in the actin-tropomyosin-myosin-MgADP complex, 3.3 Å (PDB: 8EFH) as defined by 310 AI-Copilot model.**

The binding probability scores, p(bind), for each residue in the actin-tropomyosin-myosin-MgADP complex (PDB ID: 8EFH) reflect the strength of the binding interaction at each residue position. These scores, ranging from 0 (no binding) to 1 (strong binding), were calculated using a predictive model (AI-Copilot model) that considers structural and physicochemical features of the binding sites. Higher scores denote residues with a stronger likelihood of contributing to binding interactions within the complex. The Tyrosine residues in each actin chain (B, C, D, E, F) are highlighted with yellow color.

**Supplementary Movies**

**Movie S1.** **Oxidation-induced structural changes in HMM molecules.** Representative HS-AFM movies of the non-oxidized and SIN-1-treated skeletal HMM, a double-headed fragment of myosin, on the mica-APTES surface. Scan area: 150 × 75 nm^2^, 80 × 40 pixels^2^, temporal resolution: 5 frame per second, Scale bar: 30 nm. Related to Fig. 1.

**Movie S2. Representative HS-AFM movies of the skeletal HMM bound to the non-oxidized F-actin and SIN-1 treated F-actin in the presence of 2 µM ATP.** Scan area: 150 × 75 nm^2^, 80 × 40 pixels^2^, temporal resolution: 6.7 frame per second. Scale bar: 30 nm. Related to Fig. 1.

**Movie S3. G-actin molecular structure (PDB: 2ZWH) with highlighted Tyrosine residues.** The following residues shown as blue sticks with the phenyl ring in each G-actin subdomains (SD) were highlighted: Tyr 91, Tyr 362 (SD1); Tyr 53 (SD2); Tyr 166, Tyr 169, Tyr 294, Tyr 296, Tyr 297, Tyr 306 (SD3); Tyr 188, Tyr 198, Tyr 218, Tyr 240 (SD4). The tyrosine residues were identified through mass spectrometry analysis in actin filaments from patients with rheumatoid arthritis in**^2^**. Related to Fig. 3.

**Movie S4. Computational oxidative nitration (3NT) of the highlighted Tyrosine residues in G-actin (PDB: 2ZWH).** Oxidative modifications were performed in the pyMOL using pyTM plugin (v.1.2)**^3^** by adding the nitrogen dioxide to the tyrosine residue. The oxidized atoms in the residues highlighted by pink color. Related to Fig. 3.

**Movie S5.** **Surface evaluation of G-actin structure with non-modified and modified tyrosine residues.** The successive x, y-axis rotation of the non-oxidized and oxidized G-actin structures with 45^o^ step from initial position showed the local changes in the electrostatic potential energy around oxidized residues. The map scale represents the electrostatic potential energy from -5.0 kBT (red) to 5.0 kBT (blue). Related to Fig. 3 and Fig. S2.

**Movie S6. Solvation and ionization steps for molecular dynamics simulation of the G-actin (PDB: 2ZWH).** The structure was placed in a rectangular box according to the molecular dimensions calculated by VMD software**^4^** and filled with water molecules. The cyan and yellow spheres represent Na^2+^ and Cl^-^ ions, respectively for the ionic conditions of 0.15 M NaCl.

**Movie S7. Molecular dynamics simulations for the non-modified and computationally oxidized G-actins.** MD simulation performed for 100 ns at the temperature of 300 K, 0.15 M NaCl.

**Movie S8. Computational oxidative nitration (3NT) of the highlighted Tyrosine residues in F-actin (PDB: 6BNO).** Oxidative modifications were performed in the pyMOL using pyTM plugin (v.1.2)**^2^** by adding the nitrogen dioxide to the tyrosine residue shown as sphere. The oxidized atoms in the residues highlighted by pink color. Related to Fig. 6.

**Movie S9. Tyrosine residues are involved in the binding pockets** **in the actin-tropomyosin-myosin-MgADP complex (PDB: 8EFH).** AI-Copilot model (https://310.ai/) was used to predict the binding probabilities in the actin-tropomyosin-myosin-MgADP complex.
